# Hiding in Plain Sight: Novel Observations of Plant Crypsis in a Well-Known Symbiotic System of a Hyperdiverse Tropical Forest

**DOI:** 10.64898/2026.06.12.731843

**Authors:** Gonzalo Rivas-Torres, Froilan Macanilla, Selene Escobar R

**Affiliations:** Universidad San Francisco de Quito, Estacion de Biodiversidad Tiputini, Quito, Ecuador; Universidad San Francisco de Quito, Colegio de Ciencias Biológicas y Ambientales, Quito, Ecuador

**Keywords:** ant-plant mutualism, crypsis, devil’s gardens, *Duroia hirsuta*, *Palicourea alba*, herbivory, leaf litter, *Myrmelachista schumanni*, natural history

## Abstract

Devil’s gardens are among the most striking ant–plant mutualisms in Amazonian forests. In this system, the tree *Duroia hirsuta* is associated with the ant *Myrmelachista schumanni*, which actively removes neighboring vegetation and maintains nearly monodominant patches of the host plant. Despite the apparent efficiency of this system, repeated field observations at the Tiputini Biodiversity Station, Yasuní Biosphere Reserve, revealed that individuals of *Palicourea alba* recurrently occur within active devil’s gardens. *Palicourea alba* closely resembles dead plant material, exhibiting leaf morphology and coloration that strongly mimic the surrounding litter layer, and appears to be uncommon outside these gardens. To our knowledge, this “dead-leaf” masquerade has not been previously documented in this intensively studied system, making it a particularly striking and unexpected observation. To evaluate whether this masquerade facilitates persistence within devil’s gardens, we surveyed 35 gardens and recorded *P. alba* in 19 (52.8%). When present, *P. alba* covered on average 27% of plot area, while mean herbivory across sampled leaves remained low (8.6%). Generalized linear mixed models showed that *P. alba* cover decreased significantly with increasing herbivory (F = 8.09, *p =* 0.0159), whereas herbivory increased with leaf-litter cover (F = 8.73, *p =* 0.0120). Field observations further revealed that many individuals are nearly indistinguishable from dry leaf litter, suggesting a role for visual crypsis or masquerade. Together, these results indicate initially, that the persistence of *P. alba* within devil’s gardens is mediated by a multi-layered ecological filtering process. First, masquerade likely reduces detection by *M. schumanni*, allowing seedlings to escape ant-mediated removal. Second, low herbivory suggests either enemy avoidance or reduced apparency to herbivores within the simplified understory. Third, spatial heterogeneity in leaf-litter cover may create microhabitats where both ant activity and herbivore pressure are modulated, reinforcing establishment success. This system thus represents a previously undocumented mechanism in which plant–litter resemblance enables persistence within a highly structured, biotically filtered habitat, highlighting how subtle trait-mediated interactions may modulate outcomes in otherwise strongly deterministic mutualisms.

## Introduction

Tropical ecosystems harbor the greatest biological diversity on Earth (Gentry 1988; Mittelbach et al. 2007). Despite decades of research, much of this diversity remains insufficiently documented, particularly with respect to the ecological interactions among the many organisms coexisting in these highly complex systems (Tewksbury et al. 2014). Understanding such interactions is essential for explaining the processes that structure tropical communities and sustain their extraordinary levels of diversity.

An emblematic example is the Yasuní Biosphere Reserve in the Ecuadorian Amazon (Fig. 1), widely recognized as one of the most biodiverse places on Earth (Bass et al. 2010). At local scales, Yasuní has been reported to contain more than 655 tree species and well over 900 vascular plant species per hectare in terra firme forest (Balslev et al. 1998; Valencia et al. 2004; Bass et al. 2010), and broader estimates emphasize exceptionally high arthropod richness as well (Erwin 2004). This extraordinary diversity has motivated numerous studies focused on species discovery, estimating species richness and abundance using mid and large-scale designs, quantitative analyses, genetic tools, and increasingly sophisticated monitoring technologies (Jung et al. 2026, Guevara et al. 2022, Valencia et al. 2004).

**Figure 1.**
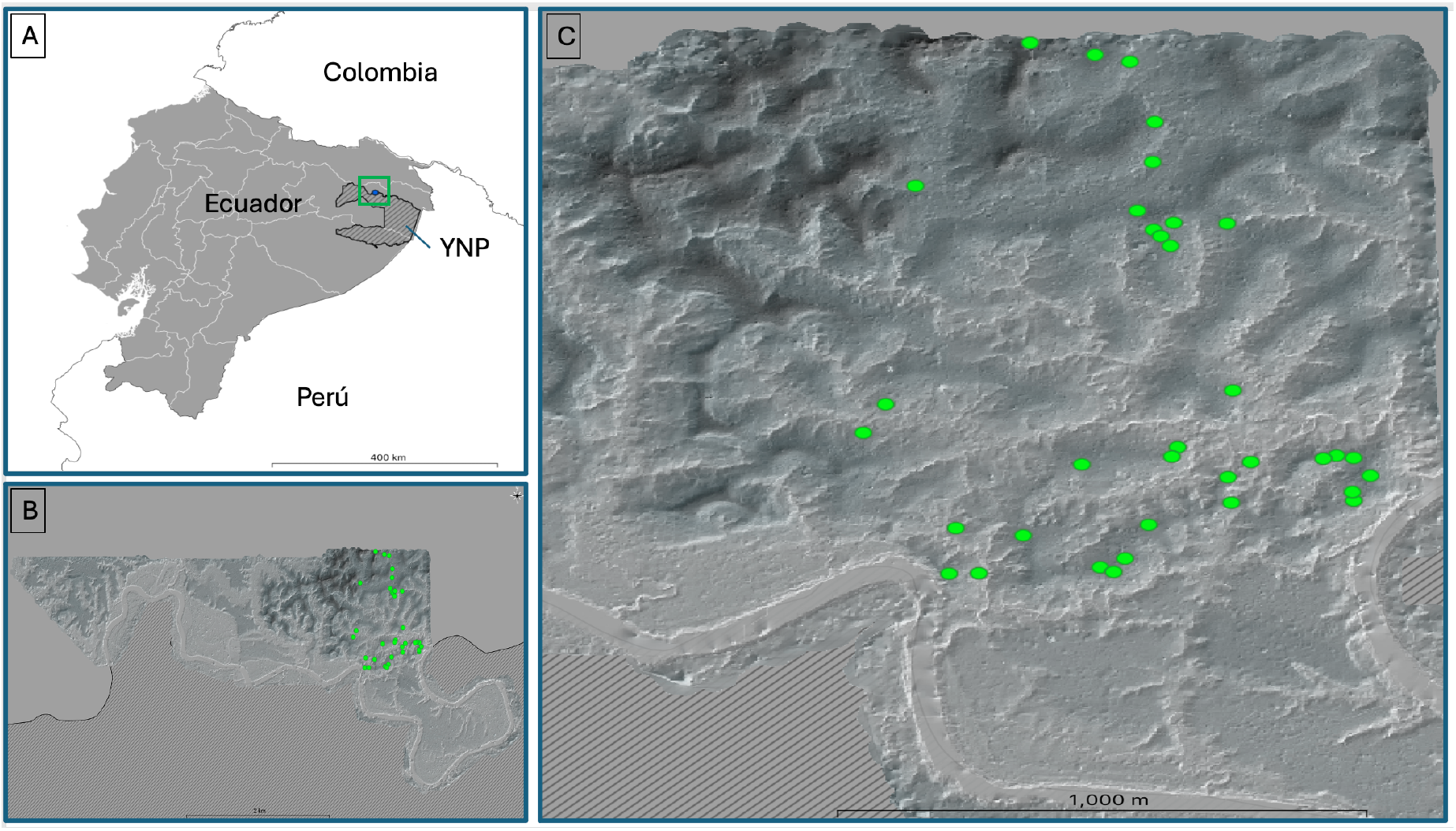
(A) Map of *Yasuní National Park* (YNP), indicated with diagonal lines, showing the location of the *Tiputini Biodiversity Station* (TBS) to the north, highlighted by a green box. (B) Digital elevation model of the 744 hectares protected by TBS. Green points indicate the location of the surveyed “devil’s gardens,” situated to the east of the station. (C) Detailed view of the location of the 35 surveyed “devil’s gardens,” randomly distributed within the area east of TBS.

Although these approaches are essential for characterizing tropical biodiversity, their predominance can overshadow other equally important lines of inquiry, such as natural history observations and the careful documentation of specific ecological interactions among species (Greene 2005, Tewksbury et al. 2014). Such observations remain crucial for revealing biological mechanisms that often go unnoticed in large-scale studies.

One of the most remarkable interactions described in Amazonian forests is the mutualistic association between the tree *Duroia hirsuta* (Rubiaceae) and the ant *Myrmelachista schumanni* (Formicidae: Formicinae). In this system, ants inhabit specialized cavities in the plant (myrmecodomatia) and contribute to the suppression of surrounding vegetation, favoring the formation of nearly monodominant stands of *D. hirsuta* known as devil’s gardens (Fig. 2) (Frederickson et al. 2005, Pfannes & Baier 2002, Cárdenas et al. 2024).

**Figure 2.**
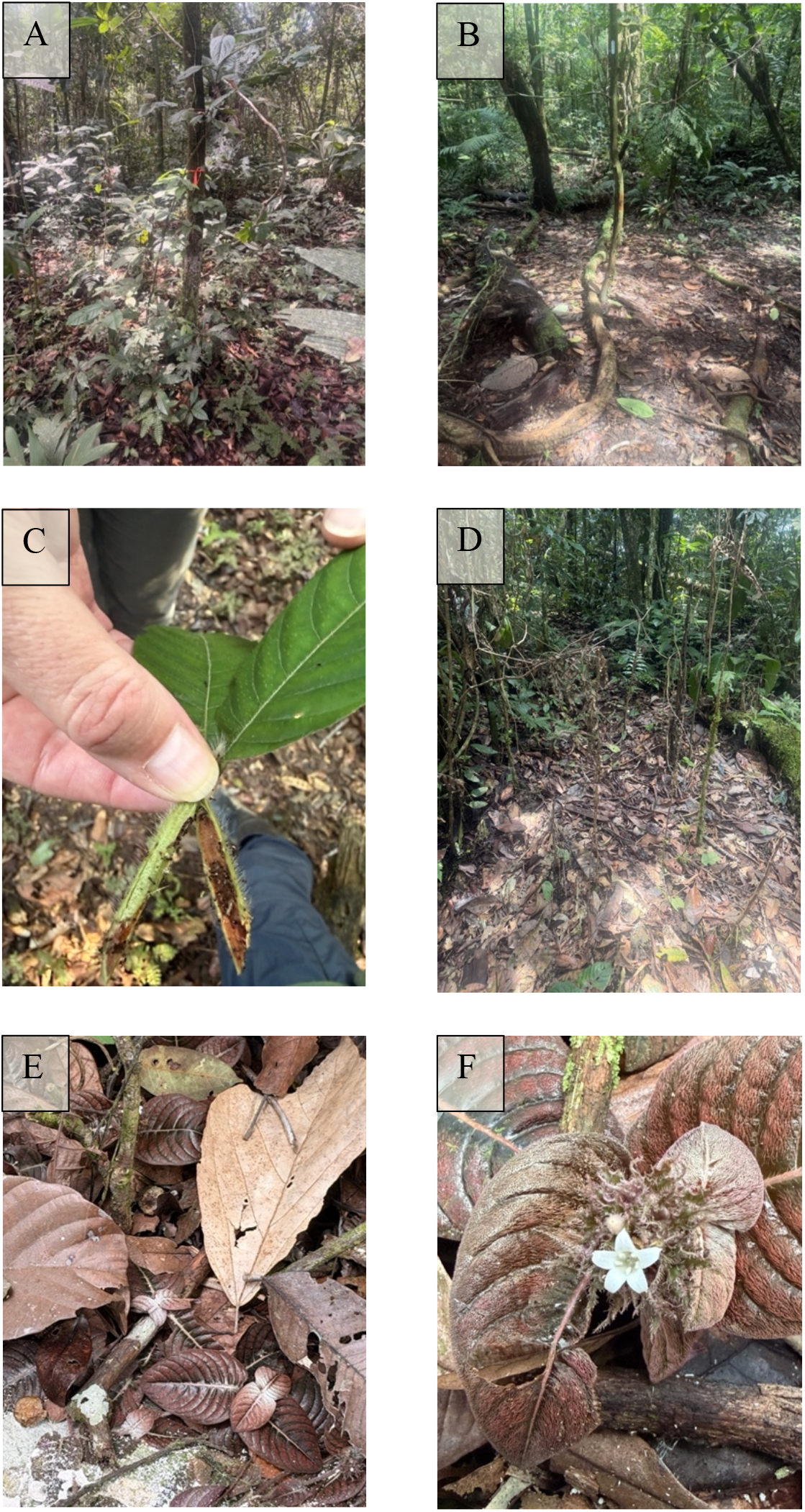
A. (A) A *Duroia hirsuta* tree without *Myrmelachista schumanni*, clearly showing the surrounding vegetation that is not eliminated by the ants. (B) A *D. hirsuta* tree at the center hosting *M. schumanni*, exhibiting a remarkably clean understory devoid of other plants. (C) Myrmecodomatia of *D. hirsuta*, where colonies of *M. schumanni* reside. (D) Mortality of seedlings and saplings of species (standing dead) other than *D. hirsuta* following attack by *M. schumanni*. (E) The striking camouflage of *Palicourea alba* at the center and lower part of the image, illustrating its masquerade among leaf litter within a plot dominated by *D. hirsuta* and patrolled by *M. schumanni*. (F) Close-up of a *P. alba* leaf showing its minute flowers and detailed leaf mimicry resembling dead leaves. In these last two panels, the low levels of herbivory experienced by *P. alba* are clearly evident.

Seminal and historical work (Pfannes and Baier 2002, Schultes 1987, Schumann 1889, Ule 1906) have shown that these gardens form because ants actively eliminate heterospecific plants by applying formic acid to their tissues, causing necrosis and death of competing seedlings (Fig. 2) (Frederickson et al. 2005). This behavior promotes the establishment and growth of *D. hirsuta* while increasing nesting opportunities for ant colonies, resulting in a highly specialized mutualism (Frederickson & Gordon 2007, Cárdenas et al. 2024).

A particularly intriguing aspect of this system is how ants distinguish their host plant from the many other species present in the understory. It has been proposed that ants use chemical signals on leaf surfaces, especially cuticular hydrocarbon profiles, as species-specific signatures. By learning the chemical profile of their host, ants may detect seedlings with different profiles during territorial patrols and trigger aggressive behavior against potential competitors (Edwards et al. 2009).

This process of chemical recognition could explain the selective elimination of non-host plants and thereby help maintain patches dominated by *D. hirsuta*, which may persist for centuries and support extremely large ant colonies. As a result, devil’s gardens are usually characterized by understory areas that are nearly free of vegetation, where most competing plants have been removed (Fig. 2) (Frederickson et al. 2005, Pfannes & Baier 2002).

Despite major advances in the understanding of this system, important questions remain about the precise mechanisms that allow ants to identify and selectively eliminate competing plants. Although chemical recognition has been proposed as a plausible explanation, it is still unclear whether this mechanism alone can explain the remarkable efficiency with which ants maintain these nearly monodominant patches. Other factors - including visual cues, plant physiological state, or temporal variation in chemical signals - may also influence ant detection and response to potential competitors (Edwards et al. 2009).

To explore this system further, we conducted several hundred hours of direct field observation in devil’s gardens at the Tiputini Biodiversity Station in the Yasuní Biosphere Reserve. These observations are part of an ongoing project aimed at documenting poorly known ecological interactions within this striking system. Here we report a novel natural history observation suggesting that plants other than *D. hirsuta* may evade attack by the associated ants. In particular, we describe the recurrent occurrence of the plant *Palicourea alba* (Rubiaceae) inside active devil’s gardens and evaluate whether their persistence may be associated with visual resemblance to the surrounding leaf litter (Fig. 2), herbivory, and other local conditions within the gardens

## Methods

### Study site and plot establishment

We surveyed 35 devil’s gardens associated with *Duroia hirsuta* at the Tiputini Biodiversity Station, Ecuadorian Amazon (Fig. 1; Supplementary Material). Gardens were selected at random within the study area. Upon arrival at each garden, we established a plot to quantify vegetation structure, ant activity, leaf litter, and the presence of *Palicourea alba*. The plot area was defined as the ground surface directly covered by the canopy projection of the *D. hirsuta* individuals forming the garden, that is, the area of physical influence beneath the trees structuring each patch. Gardens were not selected based on whether *P. alba* was present or absent, in order to avoid sampling bias.

### Vegetation surveys

Within each plot, we conducted a complete census of vegetation. We counted all individuals of *D. hirsuta*, including seedlings, juveniles, and adult trees, and we also counted all individuals of other plant species present within the plot, including seedlings, juveniles, and adult plants. Based on these counts, we classified each plot into one of three categories of heterospecific plant cover: low cover (up to 10 individuals of other plant species), medium cover (11-50 individuals), and high cover (more than 50 individuals). These categories were used as a proxy for the amount of non-*Duroia* vegetation within each devil’s garden. We also noted the occurrence of other heterospecific plants when present, but our observations focused especially on the plant *P. alba*, which occurred at relatively high frequency.

### Ant nest occupancy and activity

To estimate ant nest occupancy, we randomly selected 10 *Duroia* domatia per plot from different plants and inspected each one for the presence or absence *of Myrmelachista schumanni*. Nest occupancy frequency was calculated as the percentage of inspected domatia occupied by ants. In addition, we visually assessed ant activity within each plot by recording whether ants were actively patrolling the ground surface and the branches of *Duroia* individuals. During visits we also qualitatively documented the presence and activity of ants coming from *Duroia* trees, on vegetation and soil to verify that sampled patches corresponded to active devil’s gardens maintained by the previously described mutualism.

### *Palicourea alba* occurrence, cover, herbivory, and occurrence outside gardens

Within each plot, we recorded whether *P. alba* was present. When it occurred, we visually estimated the percentage of plot area covered by *P. alba*. To estimate herbivory, we randomly selected 10 leaves from different *P. alba* individuals in each plot and visually estimated the percentage of leaf area removed or damaged by herbivores. The 10 leaves could belong to 10 different individuals in most cases, but when lower numbers of *P. alba* were recorded, analyzed leaves could be repeated in the same individual. These values were averaged to obtain a mean herbivory estimate per plot. To evaluate whether *P. alba* was restricted to devil’s gardens or also present in surrounding areas, we searched both within each plot and within a 2-m buffer around the plot and recorded presence or absence outside the garden area.

### Leaf litter and specimen collection

We visually estimated the percentage of plot area covered by leaf litter because litter may influence both the detectability of plants and the visual background against which *P. alba* individuals occur (Skelhorn et al. 2010). During the study, we also collected voucher material of *D. hirsuta, P. alba*, and the ants present in the gardens to confirm taxonomic identities. Plant samples were collected directly from individuals observed within the gardens, and ants were collected manually from vegetation and the substrate during periods of activity. These samples were later examined to confirm the identities of the plants and associated ant colonies. Voucher ant specimens were deposited in the invertebrate collection of the Zoology Museum at Universidad San Francisco de Quito (ZSFQ) and identified using diagnostic morphological characters, supported by taxonomic literature (Longino 2006) and reference images from AntWeb (Type: CASENT0905165). Vouchers of P. alba were analyzed by G R-T and identified using virtual Herbarium material from sources like MOBOT, and Kew Gardens (www last accessed). All these materials will help as well, for next analyses to understand if chemical cues present or absent in these species, are explaining the presence of *P. alba* in devil’s gardens.

### Statistical analyses

We evaluated the relationship between *Palicourea alba* and multiple biotic and abiotic predictors using generalized linear mixed models (GLMMs) implemented in JMP SAS V.18. Analyses were conducted in two complementary steps, explicitly varying the dependent variable to test different components of the system. First, we modeled *P. alba* cover (% plot area; continuous, positive, and right-skewed) as the response variable. Fixed predictors included the number of *D. hirsuta* individuals per plot, leaf-litter cover (%), herbivory on *P. alba* (% leaf area removed), ant nest occupancy frequency of *Myrmelachista schumanni*, and heterospecific plant-density category. Given the distribution of the response, we fitted GLMMs with a Gamma error distribution and log link function. Plot identity was included as a random effect to account for spatial heterogeneity and non-independence among gardens. Model significance was assessed using likelihood-based F-tests, and diagnostics included inspection of residuals and overdispersion.

Second, to evaluate potential indirect pathways, we repeated the analysis using herbivory on *P. alba* as the dependent variable. The same set of predictors was initially considered, and model structure (Gamma distribution with log link; plot as random effect) was maintained for consistency. Together, this two-step GLMM framework allowed us to disentangle direct and indirect effects, revealing a mediated relationship in which leaf-litter cover influences *P. alba* cover indirectly through its effect on herbivory.

### Ongoing analyses

As part of a broader project that is still in progress, we are as well, conducting additional analyses to characterize interactions among these species in more detail. These include characterization of chemical profiles of *D. hirsuta, P. alba*, and *M. schumanni*, direct interaction experiments between ants and seedlings of different species, observations of ant responses to plants in different physiological states, evaluations of colony patrolling behavior, and comparative analyses of plant composition inside and outside devil’s gardens. These results will be presented elsewhere.

## Results

We found *Palicourea alba* in 19 of the 35 devil’s gardens surveyed, indicating that the species was present in more than half (52.8%) of sampled gardens. When *P. alba* was present, it covered on average 27% (±SD 23.65) of plot area across the 19 gardens in which it occurred. Mean herbivory on *P. alba* was relatively low: averaging across 10 leaves from different individuals per plot and then across the 19 occupied plots, estimated leaf damage was 8.6% (±SD 7.89).

Direct field observations repeatedly showed *P. alba* individuals growing on the leaf-litter-covered substrate of active devil’s gardens. In many cases, plants were detected only after careful inspection because their leaves closely resembled the surrounding dry litter in color, position, and general appearance (Fig. 2). In all gardens where we examined the ants associated with patches containing *P. alba*, the active colonies corresponded to *Myrmelachista schumanni*, confirming that these plants occurred within gardens maintained by the ant species previously reported for the *Duroia hirsuta*-*Myrmelachista schumanni* mutualism. During our observations, we did not record obvious recent damage (i.e. plants that look killed by formic acid) inflicted by ants on the *P. alba* individuals present inside the gardens.

### Associations with *Palicourea alba* cover

Simple relationships were explored between *Palicourea alba* cover and the number of *Duroia* individuals in plots, leaf-litter cover, herbivory, ant nest occupancy frequency, and heterospecific plant-density category. Although several of these relationships suggested marked tendencies, generalized linear mixed models with a Gamma distribution showed that only herbivory was significantly associated with *P. alba* cover (F = 8.09, *p* = 0.0159). Specifically, *P. alba* cover declined as herbivory increased, indicating that plots with higher herbivory had lower *P. alba* cover.

### Associations with herbivory

We then repeated the analysis using herbivory on *Palicourea alba* as the dependent variable. In this model, only leaf-litter cover was significant (F = 8.73, *p =* 0.0120). Based on the results from GLMMs, the relationship was positive: herbivory on *P. alba* increased as leaf-litter cover increased.

Simple relationships were explored between *Palicourea alba* cover and the number of *Duroia* individuals in plots, leaf-litter cover, herbivory, ant nest occupancy frequency, and heterospecific plant-density category. Although several of these relationships suggested marked tendencies, generalized linear mixed models with a Gamma distribution showed that only herbivory was significantly associated with *P. alba* cover (F = 8.09, *p* = 0.0159). Specifically, *P. alba* cover declined as herbivory increased, indicating that plots with higher herbivory had lower *P. alba* cover.

## Discussion

This pattern is consistent with at least four non-mutually exclusive interpretations. First, devil’s gardens may function as enemy-free space for *P. alba*, if ant activity indirectly reduces herbivore pressure and thereby allows plants to persist and expand in cover (Jeffries & Lawton 1984). Second, herbivory may act as an additional ecological filter within devil’s gardens: even if some heterospecific plants escape ant detection, they may still be restricted by herbivores. Third, *P. alba* may be restricted to specific microhabitats within gardens where herbivore activity is lower. Fourth, if *P. alba* resembles leaf litter, that resemblance may reduce detection not only by ants but also by herbivores, so that crypsis could confer a dual advantage (Skelhorn et al. 2010).

Taken together, the two models suggest an indirect pathway: greater leaf-litter cover was associated with greater herbivory on *P. alba*, and greater herbivory was associated with lower *P. alba* cover. In conceptual terms, this indicates a mediated interaction in which leaf litter may influence *P. alba* abundance through its effects on herbivory.

This result is biologically striking because it is not the pattern expected if litter-mediated camouflage simply reduced herbivory. A straightforward expectation would be that more litter should provide better background matching and therefore lower herbivory. Instead, our data (Supplementary Material) suggest the opposite. One plausible explanation is that leaf litter provides habitat or refuge for understory herbivores, thereby increasing local herbivore abundance and damage to *P. alba*. Many potential understory herbivores occur within litter layers, including orthopterans, coleopterans, larvae, isopods, and other arthropods.

These findings therefore suggest a possible trade-off: growing in sites with greater leaf litter may improve camouflage and reduce detection by ants, while simultaneously increasing exposure to herbivores that inhabit the litter layer. Under this scenario, *P. alba* may persist only where there is enough litter to reduce ant detection but not so much litter that herbivory becomes excessive. In other words, persistence within devil’s gardens may depend on a narrow ecological window shaped by multiple interacting biotic filters.

The recurrent presence of *Palicourea alba* inside devil’s gardens is particularly interesting because these patches are maintained by two ecological processes generally considered highly effective: the direct elimination of competing plants by *Myrmelachista schumanni* and the possible allelopathic effects associated with *Duroia hirsuta* (Frederickson et al. 2005, Page et al. 1994). In this context, the persistence of *P. alba* suggests that at least some plant species may establish within these patches through alternative routes that have not been documented previously.

If visual resemblance to leaf litter effectively reduces detection by ants, this system may represent a poorly documented example of plant crypsis or masquerade in a plant-insect interaction (Skelhorn et al. 2010). Such mechanism would imply that morphological or phenotypic traits associated with leaf appearance influence the probability that a plant is detected or attacked by the ants maintaining patches dominated by *D. hirsuta*.

At the same time, our quantitative results indicate that persistence cannot be explained solely by putative camouflage. *P. alba* cover decreased with herbivory, yet herbivory increased with leaf-litter cover. Considered together, these patterns suggest that *P. alba* may be shaped by opposing ecological pressures. Leaf litter may enhance visual crypsis and reduce detection by ants if any, but it may also increase herbivore activity, leading to higher herbivory. This trade-off may restrict *P. alba* to a narrow range of local conditions within devil’s gardens (Jeffries & Lawton 1984, Skelhorn et al. 2010).

These observations raise several mechanistic questions. It will be important to determine whether the apparent evasion by *P. alba* is driven primarily by foliar visual traits or whether other factors also contribute, such as changes in leaf chemistry, particular physiological states, or variation in ant patrolling intensity (Edwards et al. 2009). The relatively high frequency with which we found *P. alba* within gardens suggests that this phenomenon may not be exceptional and raises the possibility that other plant species may also use comparable strategies to persist within these patches.

If confirmed, this pattern may also have broader evolutionary implications for understanding how plants persist in environments with especially strong biotic pressure. Devil’s gardens represent a system in which competing plants face simultaneous pressure from ant-mediated removal and possible host-associated chemical effects. In that setting, resemblance of *P. alba* to surrounding litter could represent an adaptive trait that reduces the probability of detection or attack, allowing persistence where most other heterospecific species are eliminated (Frederickson et al. 2005, Skelhorn et al. 2010).

As next steps that are part of on-going efforts, it will be necessary to evaluate experimentally the response of *M. schumanni* to *P. alba* individuals and to plants with manipulated visual or physiological states. Manipulation of foliar appearance, comparative analyses of chemical profiles of *P. alba* and *Duroia*, and quantitative assessment of the abundance of *P. alba* in relation to ant activity and litter structure should help distinguish among visual, chemical, and combined mechanisms. We are also evaluating whether other ant species may occupy *Duroia* individuals and whether shifts in occupancy or reduced activity of *M. schumanni* may correlate with the occurrence of *P. alba*. Preliminary observations suggest that some *Duroia* trees may show lower ant activity or lower frequencies of occupancy, raising the possibility that variation in ant presence or species composition could influence plant structure inside the gardens.

### Natural history significance and broader relevance

Detailed natural history observations have historically been fundamental to the development of ecology, because many central hypotheses and theories first emerged from careful field documentation of biological interactions. In recent decades, however, increasing emphasis on macroecological approaches, large-scale analyses, and complex analytical tools has relatively reduced attention to direct observation of organisms in their natural environments. Yet natural history remains a critical source of discovery, especially in tropical ecosystems where most species interactions remain poorly known (Greene 2005, Tewksbury et al. 2014). Our case illustrates this point clearly. Although devil’s gardens associated with *Duroia hirsuta* and *Myrmelachista schumanni* have been the subject of ecological and evolutionary study for decades, the observations presented here suggest that even systems considered relatively well known may still harbor undocumented interactions. This finding reinforces the idea that complete understanding of ecological systems requires complementing large-scale and analytical approaches with repeated, prolonged, direct field observation (Pfannes & Baier 2002, Frederickson et al. 2005, Cárdenas et al. 2024, Greene 2005).

With this note, we also hope to encourage biologists, naturalists, and ecologists - especially younger researchers - to revisit systems that appear to be well understood. Even after decades of study, new observations can still reveal interactions, behaviors, or mechanisms that had previously gone unnoticed, thereby enriching our understanding of ecological and evolutionary processes (Tewksbury et al. 2014). We also intend to disseminate future results from this system through digital platforms and broader scientific outreach venues so that these findings can reach a wide audience. In highly complex tropical forests such as those of Amazonia, careful observation of organisms in their natural settings remains indispensable for understanding ecological and evolutionary dynamics and, ultimately, for informing long-term conservation and management (Greene 2005, Tewksbury et al. 2014).

## Supporting information

Supplementary Material Duroias and ants

## Acknowledgments

We want to thank the Tiputini Biodiversity Station and Universidad San Francisco de Quito for providing the necessary funding to develop this work. Special thanks to Nina Witteveen for the valuable help during part of the fieldwork for this study. We want to thank also the Ministry of Environment of Ecuador for providing permit number MAATE-ARSFC-2025-0319 necessary to perform this research.

## Notes

### Competing Interest Statement

The authors have declared no competing interest.

### Summary of Updates

We just need to include one line in the acknowledgements section to include Mrs Witteveen: We want to thank the Tiputini Biodiversity Station and Universidad San Francisco de Quito for providing the necessary funding to develop this work. Thanks to Nina Witteveen for the valuable help during part of the field work of this study. We want to thank also the Ministry of Environment of Ecuador for providing permit number MAATE-ARSFC-2025-0319 necessary to perform this research.

