## Supplementary Material Duroias and ants for "Hiding in Plain Sight: Novel Observations of Plant Crypsis in a Well-Known Symbiotic System of a Hyperdiverse Tropical Forest"

Plot# represents the plot number assigned in the field to each sampling unit; Plant.Cat. is the classification of each plot into one of three categories of heterospecific plant cover: low cover (up to 10 individuals of other plant species), medium cover (11–50 individuals), and high cover (more than 50 individuals); Ant.Frec. is the percentage of *Duroia* domatia occupied by *Myrmelachista schumanni*, estimated by randomly selecting 10 domatia per plot from different plants and inspecting each domatium for the presence or absence of ants; Ant.Patrol is whether ants were observed descending the stems of *Duroia* individuals and patrolling the plot floor; %Pali is the percentage cover of *Palicourea alba* within the plot; %Leaf.Litt. is the percentage of plot area covered by leaf litter; #Duroia is the number of *Duroia* individuals recorded within the plot; %Herbiv.Pali is the level of herbivory on *P. alba*, estimated by randomly selecting 10 leaves from different *P. alba* individuals per plot and visually estimating the percentage of leaf area removed or damaged by herbivores; Pali.Pres. is the presence or absence of *P. alba* within the plot; Pali.out is whether any individuals of *P. alba* were recorded outside the plot boundaries; Date is the date on which data for each plot were collected.

| Plot# | Plant.Cat. | Ant.Frec | Ant.Patrol | %Pali | %Leaf.Litt. | #Duroia | %Herbiv.Pali | Pali.Pres. | Pali.out | Date |
|---|---|---|---|---|---|---|---|---|---|---|
| 700 | HIGH | 60 | YES | NO | 90 | 15 | NO | NO | NO | March/17/2026 |
| 701 | HIGH | 50 | YES | NO | 85 | 10 | NO | NO | NO | March/17/2026 |
| 702 | LOW | 70 | YES | NO | 50 | 1 | NO | NO | NO | March/17/2026 |
| 704 | LOW | 100 | YES | 60 | 70 | 7 | 3.3 | YES | NO | March/17/2026 |
| 705 | LOW | 70 | YES | NO | 80 | 4 | NO | NO | NO | March/17/2026 |
| 706 | LOW | 100 | YES | NO | 70 | 1 | NO | NO | NO | March/17/2026 |
| 708 | LOW | 100 | YES | NO | 90 | 8 | NO | NO | NO | March/17/2026 |
| 709 | LOW | 100 | YES | NO | 80 | 2 | NO | NO | NO | March/17/2026 |
| 710 | LOW | 100 | YES | NO | 95 | 34 | NO | NO | NO | March/17/2026 |
| 711 | LOW | 60 | YES | NO | 80 | 22 | NO | NO | NO | March/17/2026 |
| 712 | LOW | 100 | YES | NO | 90 | 1 | NO | NO | NO | March/17/2026 |
| 713 | LOW | 100 | YES | NO | 70 | 1 | NO | NO | NO | March/17/2026 |
| 714 | LOW | 80 | YES | NO | 90 | 14 | NO | NO | NO | March/17/2026 |
| 715 | MID | 100 | YES | 80 | 50 | 4 | 5.5 | YES | YES | March/18/2026 |
| 716 | HIGH | 100 | YES | NO | 70 | 18 | NO | NO | NO | March/18/2026 |
| 717 | LOW | 100 | YES | NO | 95 | 7 | NO | NO | NO | March/18/2026 |
| 718 | HIGH | 90 | YES | NO | 80 | 10 | NO | NO | NO | March/18/2026 |
| 719 | HIGH | 80 | YES | 10 | 90 | 30 | 6.5 | YES | NO | March/18/2026 |
| 720 | MID | 90 | YES | 8 | 80 | 1 | 10.7 | YES | NO | March/18/2026 |
| 721 | LOW | 90 | YES | 50 | 60 | 2 | 3.2 | YES | NO | March/18/2026 |
| 722 | MID | 100 | YES | 2 | 70 | 10 | 21 | YES | NO | March/18/2026 |
| 723 | MID | 100 | YES | 40 | 70 | 40 | 2.2 | YES | YES | March/18/2026 |
| 724 | MID | 100 | YES | 30 | 70 | 3 | 2.4 | YES | NO | March/18/2026 |
| 725 | LOW | 100 | YES | 30 | 80 | 5 | 12.2 | YES | NO | March/18/2026 |
| 726 | LOW | 100 | YES | 1 | 90 | 16 | 28.1 | YES | NO | March/18/2026 |
| 727 | LOW | 100 | YES | 70 | 60 | 11 | 1.3 | YES | NO | March/18/2026 |
| 728 | LOW | 100 | YES | 70 | 20 | 12 | 2.1 | YES | NO | March/18/2026 |
| 729 | LOW | 100 | YES | 2 | 80 | 36 | 5.3 | YES | NO | March/18/2026 |
| 730 | LOW | 90 | YES | 3 | 85 | 5 | 7.5 | YES | NO | March/18/2026 |
| 731 | MID | 100 | NO | 30 | 90 | 7 | 12.8 | YES | NO | March/18/2026 |
| 732 | LOW | 100 | YES | 10 | 60 | 13 | 0.9 | YES | NO | March/18/2026 |
| 733 | LOW | 100 | YES | 7 | 80 | 36 | 17.5 | YES | NO | March/18/2026 |
| 734 | HIGH | 100 | NO | 1 | 85 | 11 | 16 | YES | NO | March/18/2026 |
| 735 | LOW | 100 | NO | 5 | 90 | 9 | 4.5 | YES | NO | March/18/2026 |
| 736 | LOW | 100 | YES | NO | 90 | 1 | NO | NO | NO | March/18/2026 |
